# Phenotypic characterization of *Prevotella melaninogenica ATCC 25845*

**DOI:** 10.64898/2026.09.25.754359

**Authors:** Claire Albright, Sophie M. P. Pickelner, Souzane Ntamubano, Gouri Anil, Kelyah Spurgeon, Ariangela J. Kozik

## Abstract

*Prevotella melaninogenica* is a Gram-negative bacterium that has been implicated in numerous disorders such as chronic obstructive pulmonary disease (COPD) and cystic fibrosis, and despite its abundance in the human microbiome, there is still limited knowledge on the fundamental physiology describing *Prevotella melaninogenica*. To properly advance *Prevotella* biology, it is critical to report and document such characterizations of *Prevotella;* therefore, the purpose of this study is to examine and define the phenotypic and growth characteristics of *P. melaninogenica* ATCC 25845. In this study, we found that *Prevotella melaninogenica* ATCC 25845 is a Gram-negative bacterium that is predominantly short and medium rod-shaped, with occasional filamentous rods. SEM imaging revealed that this filamentation may serve as division centers, points from which multiple daughter cells form. We also describe the antibiotic profile, which showed susceptibility to four classes and resistance to six classes of antibiotics. Carbon source growth kinetic characterization revealed a preference for carbohydrate sources.

**Importance:** The growth characteristics of *P. melaninogenica* ATCC 25845 are poorly understood. No formal studies have investigated its fundamental physiological characteristics. This study defines the phenotypic and growth characteristics of *P. melaninogenica* ATCC 25845 by exploring morphology and replication phenomena not previously described. Additionally, our findings deepen understanding of *P. melaninogenica* behavior across diverse environments. Conducting a comprehensive carbon utilization panel of over 90 carbon sources and antibiotics, we provide a high-throughput analysis of the physiology of *P. melaninogenica* ATCC 25845.

## Introduction

*Prevotella melaninogenica* is a non-motile, non-sporulating, Gram-negative bacterium (1). While the organism was first described by Oliver and Wherry in 1921 as *Bacteroides melaninogenicus* (2), it was later reclassified and split into the *Prevotella* genus as *Prevotella melaninogenica* by Shah and Collins, where it has since remained despite extensive taxonomic reorganization within the genus (3-4). Its taxonomic stability, however, contrasts with its historically limited understanding of its ecological significance. Although members of the *Prevotella* genus have long been recognized as members of the human microbiome, the true abundance and prevalence of *P. melaninogenica* remained severely underappreciated. Traditional culture-based methods were restricted by the organism’s oxygen sensitivity and fastidious growth requirements, and early cultivation-independent approaches lacked species-level identification. Nevertheless, recent advances in metagenomics and whole-genome sequencing have overcome this limitation, enabling culture-independent, species-level resolution within the respiratory microbial communities (5-6). Therefore, we now understand that *P. melaninogenica* is not merely present but is a significant member of the human respiratory tract, accounting for approximately 4–8% of the oral microbiome and roughly 10% of the lower airway microbiome in healthy individuals (7-8).

While *P. melaninogenica* abundance is associated with respiratory health and disease states, its role and function within respiratory ecosystems remain poorly understood. Research has implicated *P. melaninogenica* in maintaining respiratory health (9-10), yet the organism is also associated with a broad spectrum of respiratory disorders, including chronic obstructive pulmonary disease (COPD), cystic fibrosis, gastroesophageal reflux disease, and long COVID (11-15). Collectively, these findings suggest that *P. melaninogenica* influences both health and disease states, potentially in a context-dependent manner. Despite its clinical relevance, the core physiology and metabolic profile of this bacterium remain largely uncharacterized. This knowledge gap is primarily attributable to the technical challenges associated with culturing this fastidious organism. Consequently, research has largely stalled at the association level, impeding direct investigations into how *P. melaninogenica* modulates the respiratory environment. Nevertheless, the successful colonization of *P. melaninogenica* across strikingly varied environments underscores its relevance to microbiome community dynamics and the human host (16). Therefore, this study provides the first detailed characterization of the phenotypic and metabolic profile of *P. melaninogenica*, defining the fundamental physiological traits of this highly abundant member of the human respiratory tract.

The *P. melaninogenica* strain examined in this study is ATCC 25845. We chose this strain because it is the type strain, and the limited existing data on *P. melaninogenica* stems primarily from studies conducted on ATCC 25845 (1, 17-18). Strain ATCC 25845 was originally recovered from sputum at the Wadsworth Anaerobic Laboratory and was later deposited in the American Type Culture Collection (ATCC) by the Virginia Polytechnic Institute (1). Although previous work has provided valuable insight into *Prevotella* biology, the molecular microbiology and physiology of *P. melaninogenica* remain underexplored, and comprehensive, updated phenotypic data are limited. The present study addresses this gap by defining the phenotypic and growth characteristics of *P. melaninogenica* ATCC 25845.

## Results

### P. melaninogenica ATCC 25845 is a Gram-negative coccobacillus with pigment formation capabilities

Consistent with previous findings (19), our data suggest that *P. melaninogenica* ATCC 25845 is a Gram-negative coccobacillus. Grown under strictly anaerobic conditions (0% O2, 5% H2, and 95% N2) at 37°C, the cellular and colony morphologies remained largely consistent across a variety of commercially available and lab-manufactured nutrient and selective agar, with slight phenotypic differences in media containing sheep’s blood as a principal component (Fig. 2). After 48 hours of anaerobic incubation, *P. melaninogenica* ATCC 25845 exhibited a distinct tan pigment with a cream center. Pigment production was only observed on Brucella Blood Agar (BRU), Biolog Universal Growth Agar (BUA), Laked Brucella Blood Agar w/ Kanamycin and Vancomycin (LKV), and *Prevotella* Selective Agar (PSA). On media without sheep’s blood, such as Brain Heart Infusion (BHI) and supplemented Tryptic Soy Agar (TSA+), colonies appeared opaque and cream-colored. Nevertheless, on all agar media tested, the colonies were round, with entire edges and raised elevation. Microscopic images showed minimal variation between the agar and broth media tested. Gram stain images revealed Gram-negative staining with predominantly short and medium rod-shaped cells with occasional filamentous rods (Fig. 1). These observations provide a detailed characterization of the diverse visual morphologies observed in *P. melaninogenica* ATCC 25845, notably pigment production on solid media containing sheep’s blood.

**Figure 1.**
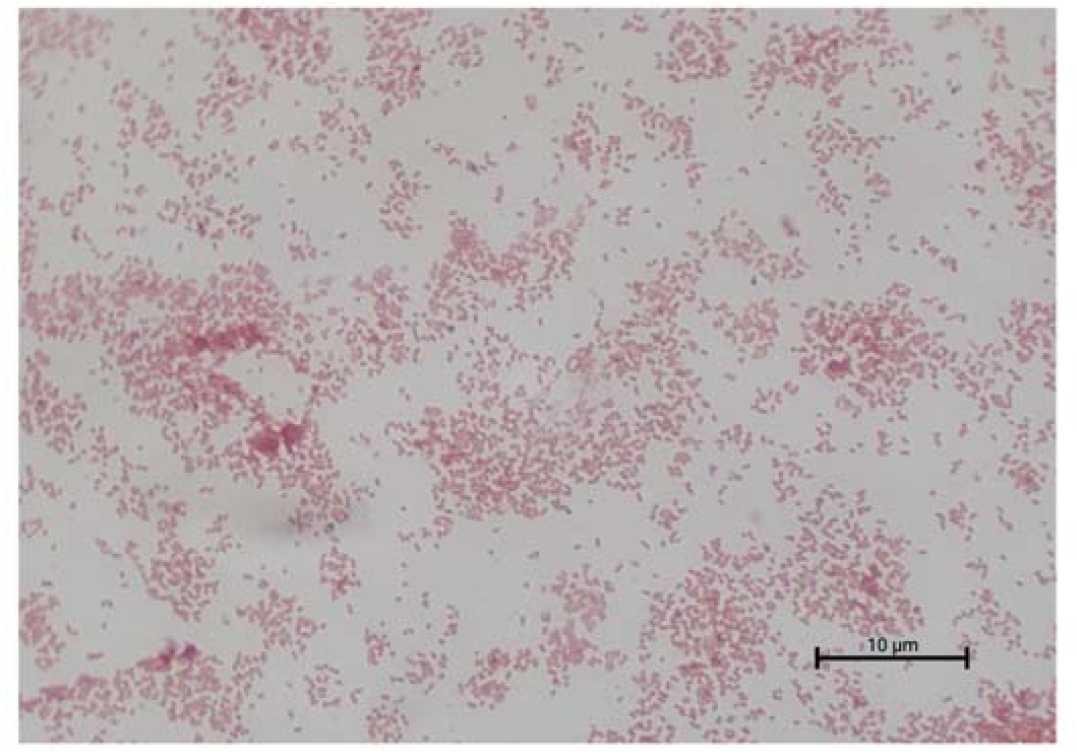
Prevotella melaninogenica 25845 morphology. Gram stain of *Prevotella melaninogenica* displaying Gram-negative coccobacillus morphology. The organism was cultured on solid brain heart infusion (BHI) media for 24 h under anaerobic conditions prior t staining and imaging. Scale bar = 10μm.

**Figure 2.**
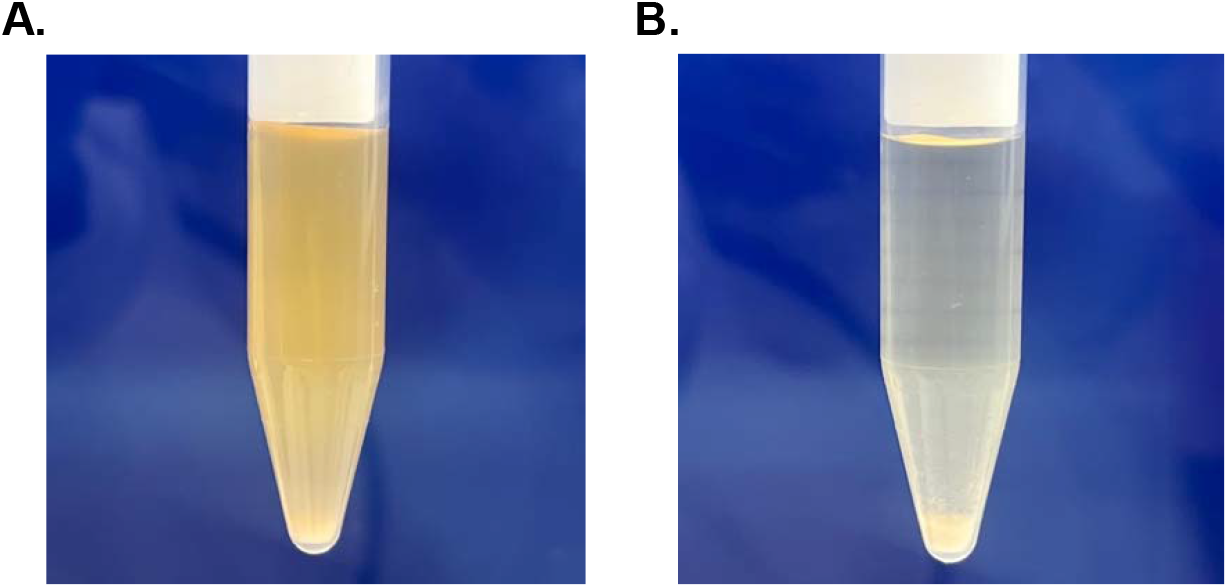
*Prevotella melaninogenica* culture morphology varies depending on the liquid media. Liquid culture phenotypes of *P. melaninogenica* ATCC 25845. **(A)** Growth in liquid BHI broth shows partial homogeneity with minor sedimentation. **(B)** Growth in liquid TSA broth exhibits near-total cell sedimentation. Cultures (n= 3 biological replicates per condition) were grown for 24 hrs at 37ºC under anaerobic conditions prior to imaging. Images are representative of phenotypes observed consistently across all biological replicates and batches.

### Cell size of P. melaninogenica decreases along with growth phase progression

We aimed to define the growth kinetics and cellular morphology of *P. melaninogenica* over time under anaerobic conditions. To achieve this objective, the inoculating culture of *P. melaninogenica* was initially grown in BHI broth at 37°C under strictly anaerobic conditions. Inoculums derived from this culture were then cultured in BHI broth at 37°C. Growth was quantified by measuring optical density (OD600) over 24 hours, and the number of viable cells was determined by anaerobic plating to quantify colony-forming units (CFU). The identity of the bacteria was confirmed using Gram stain and PCR. A negative control consisting of cell-free media was measured in parallel. To assess phase-dependent morphology, culture samples were collected anaerobically during the lag, exponential, and stationary phases of growth for subsequent analysis via scanning electron microscopy (SEM). The resulting growth curve demonstrated that *P. melaninogenica* has a replication rate of 1.3 hours, reaching a maximum optical density of ∼0.33 over 12 hours. During the stationary phase, a slight decrease in optical density to ∼0.28 was observed (Fig. 3A). Anaerobic CFU plating at hour 0 demonstrates a viable cell count of 10^6^ CFU/ml at the time of inoculation (Fig. 3C). Subsequent anaerobic plating at hour 12 shows a significant increase to 10^9^ CFU/ml by the peak optical density, confirming robust cell division during the exponential phase. Further supporting our growth curve observations, the 24-hour anaerobic plating shows a decrease in viable cells to 10^8^ CFU/ml during the stationary phase, suggesting that the decrease in OD observed may be due to cell lysis (Fig. 3A). SEM imaging of samples collected over time revealed distinct morphological differences between growth phases. Cells in the lag and exponential phases were primarily short-to-medium rods, though both groups showed long filamentous outliers. By contrast, stationary phase populations consisted entirely of very short coccobacilli, with no filamentous cells present. Quantification of the difference in cell length between the exponential and stationary phases yielded a highly significant result (p<0.0001; Fig. 3B). These data define the replication rate of *P. melaninogenica* and demonstrate that the stationary phase is characterized by both a decrease in culture viability and a significant transition in cellular morphology from rod-shaped to coccobacilli.

**Figure 3.**
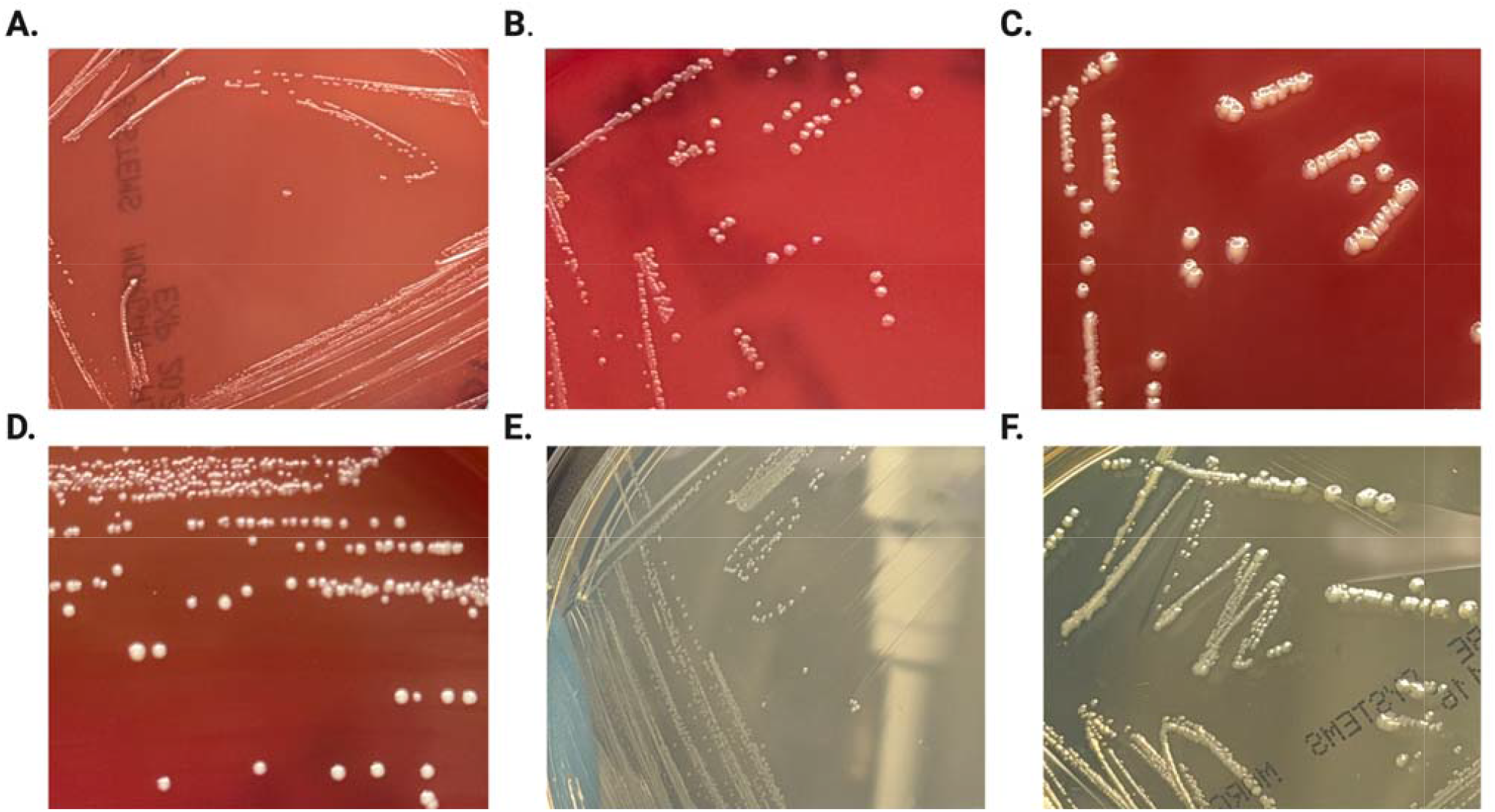
*Prevotella melaninogenica* colony morphology varies depending on the solid media. Colony morphology of *P. melaninogenica* ATCC 25845 on nutrient and selective agar media. Pigmente colony morphology was observed on LKV **(A)**, BRU **(B)**, PSA **(C)**, and BUA **(D)**.Opaque cream-colored colon morphology was observed on TSA+ **(E)** and BHI **(F)**. The observed phenotypes were reproducible across independent experimental batches.

### P. melaninogenica achieves striking filamentation before coccobacilli formation

We sought to identify any novel characteristics of growth and replication revealed by the high-resolution SEM images captured across *P. melaninogenica* growth phases. To do this, we examined lag, exponential, and stationary phase images for outliers in morphology. This analysis revealed that within the lag and exponential phases, some cells were extremely filamentous, with the longest cell in the exponential phase measuring a staggering ∼100 µm (Fig. 4A). Compared to the mean exponential phase cell length of 2µm, this represents a striking change in morphology. In addition to the filamentous rods, we detected a second notable observation: many of the filamentous cells appear to undergo division at multiple points simultaneously, resulting in multiple daughter cells instead of the classical binary fission. In a Gram stain of *P. melaninogenica* grown in BHI broth, many small rods appear lined up, as if they had just undergone simultaneous division from one large filamentous cell. This observation is supported by SEM images demonstrating filamentous cells in lag and exponential phase with multiple septum-pinching sites (Fig. 4B). These findings suggest that *P. melaninogenica* may present a novel mechanism of growth and replication previously unknown within *Prevotella* and other closely related genera.

**Figure 4.**
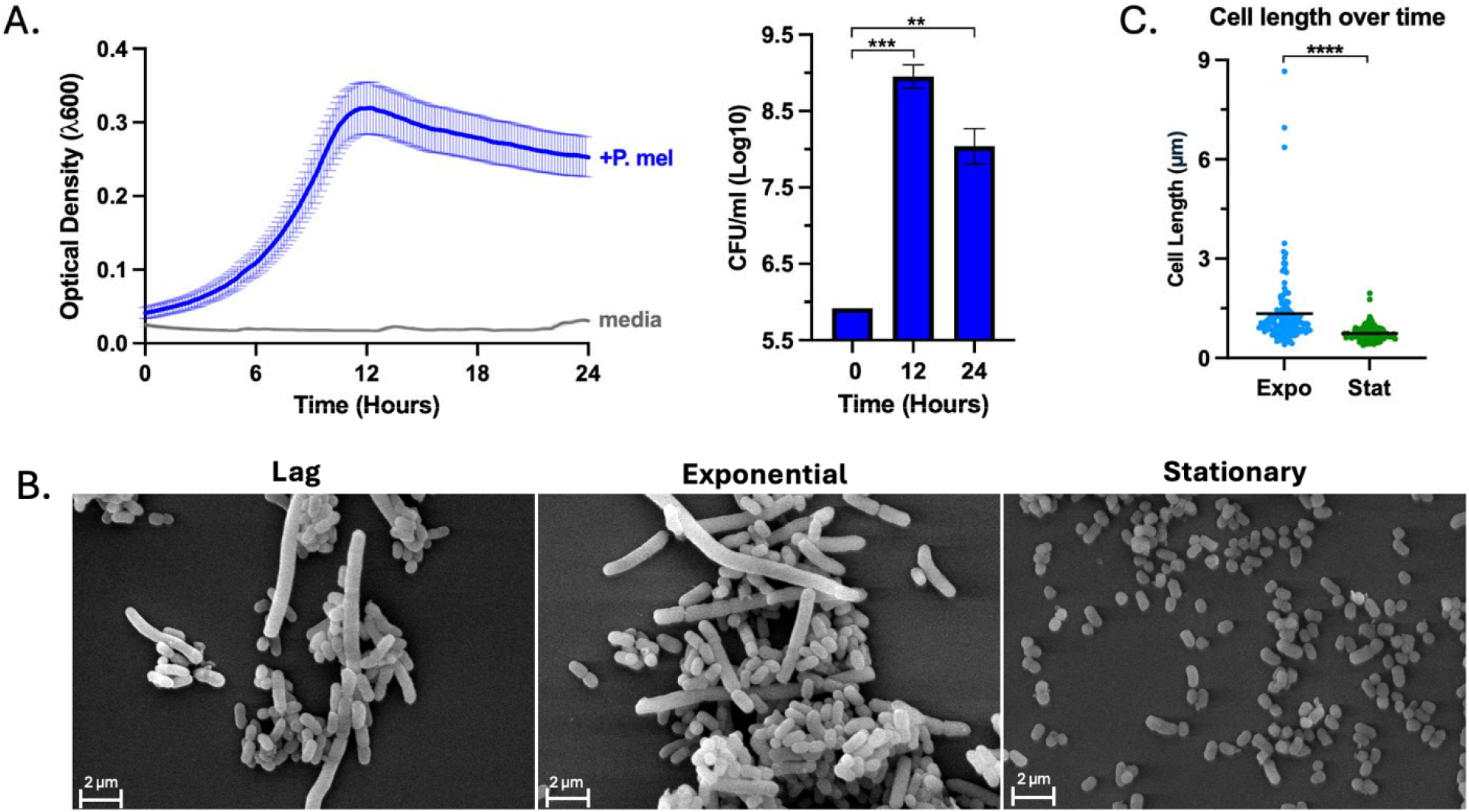
Growth curve kinetics & associated changes in cell morphology. *P. melaninogenica* ATCC 25845 exhibits a 3.21-hour replication rate (*r.r.*) and significant growth phase- dependent changes in cell morphology. **(A)** Culture optical density () was recorded over 24 hours (n = biological replicates). BHI media + *P. mel* (Blue line); BHI media – *P. mel* (Grey line). Viable cell densit (CFU/mL) was quantified at 0, 12, and 24 hrs under anaerobic conditions (n = 2 technical plating replicates per biological replicate). Statistical significance for differences in cell viability (ΔCFU) was determined using an unpaired Welch’s -test with a Gaussian distribution assumption. ΔCFU was statistically significant between 0–12() and 0–24 h (p = 0.004). **(B)** Scanning electron microscopy (SEM) images of cells harvested durin lag, exponential, and stationary growth phases (12,000× magnification; scale bar = 2 µm). **(C)** Quantification of cell length (µm) in exponential versus stationary phases (randomly selected cells per condition, quantified via ImageJ). Statistical significance for differences in cell length (ΔCell Length) was determined using an unpaired Welch’s -test with a Gaussian distribution assumption. ΔCell Length between exponential and stationary phases was statistically significant (p < 0.0005).

### P. melaninogenica is susceptible to four classes and resistant to six classes of antibiotics

Given the significant relationship between antibiotics and microbes, our next objective was to build an antibiotic characterization profile for *P. melaninogenica*. To establish this profile, bacterial growth was monitored in the presence of 44 antibiotics. These antibiotics, representing 15 distinct classes, were evaluated at four concentrations, using *P. melaninogenica* cultured in antibiotic-free media as a positive control. The inoculating *P. melaninogenica* culture was initially grown in TSB+ at 37°C under strictly anaerobic conditions. Subsequent TSB+ cultures were prepared from this inoculating culture and plated onto Biolog Phenotype MicroArray (PM) Microplates 11C and 12B. Along with the positive control, several additional antibiotics were run in a separate 96-well plate. Growth was quantified by measuring optical density (OD740) over 24 hours. Susceptibility categories were determined using Area Under the Curve (AUC) values for each antibiotic class across the four concentrations (Fig. 5A), placing each antibiotic in a sensitive, dose-dependent, or resistant grouping.

**Figure 5.**
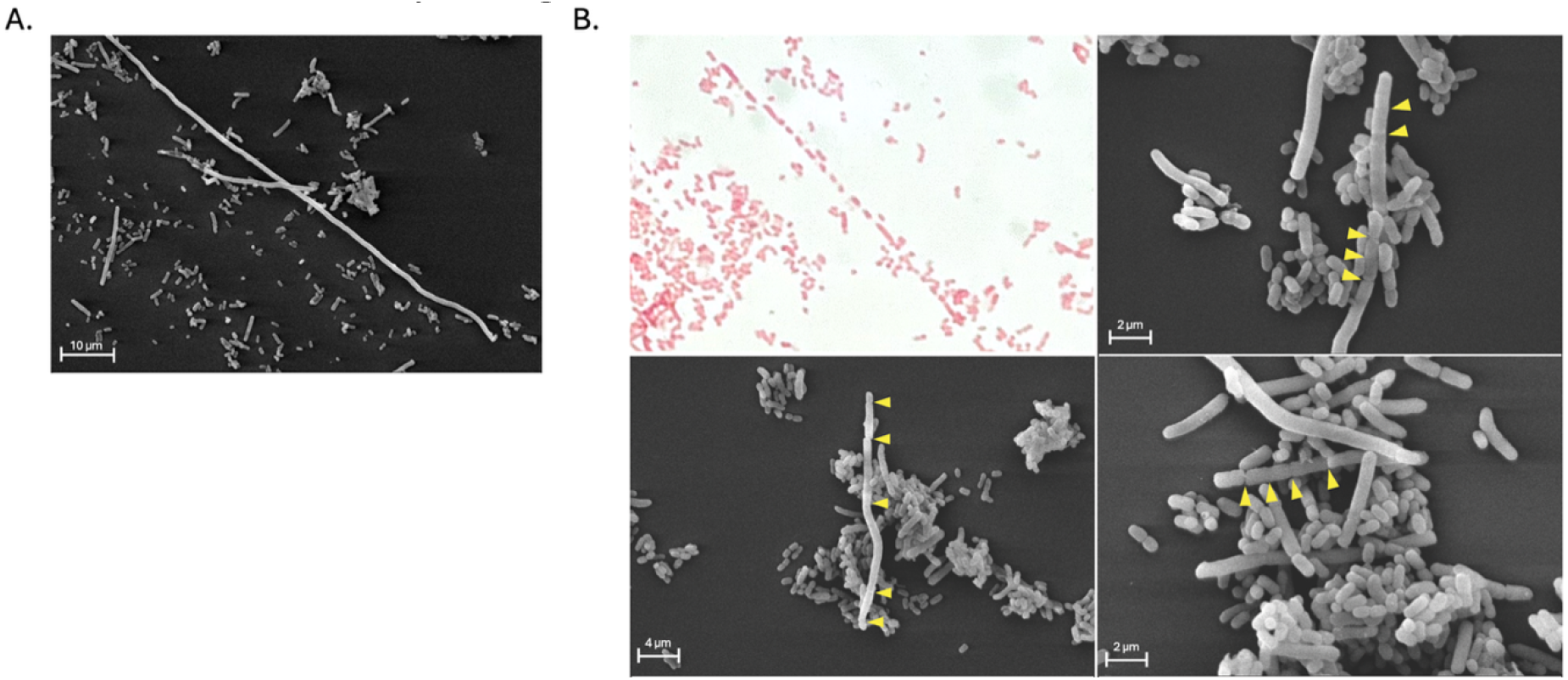
*P. melaninogenica* may become extremely filamentous and undergo spontaneous simultaneous division, yielding short rods and coccobacilli. Scanning electron microscopy (SEM) images of P. melaninogenica filamentous morphology and division during exponential growth phase. **(A)** Representative SEM image showing an elongated, filamentous cell. These filaments, with a length of ≥ 100 μm, are outliers to the short rod morphology found in exponential phase. Scale bar = 10μm. **(B)** Exponential-phase filamentous cells undergoing simultaneous multiple fission. Yellow arros denote sites of septal pinching. Scale bar = 10μm.

*P. melaninogenica* exhibited complete resistance to all tested concentrations of aminoglycoside, quinolone, glycopeptide, sulfonamide, polymyxin, and aminocyclitol antibiotics. Conversely, several classes, including oxacephem, macrolide, aminocoumarin, and lincosamide antibiotics, showed complete sensitivity, characterized by no growth over the 24-hour period. Dose-dependent growth responses were observed to varying degrees for antibiotics within the penicillin, cephalosporin, tetracycline, amphenicol, and aminocyclitol classes. While growth responses were generally consistent within most antibiotic classes, the penicillin class demonstrated significant internal variation. *P. melaninogenica* was completely sensitive to five of the seven penicillin antibiotics tested, showing no growth across all four concentrations (Supp. Fig. 1). In contrast, amoxicillin elicited a dose-dependent response, with growth observed only at the two lowest antibiotic concentrations (Fig. 6). A similar dose-dependent pattern was observed for carbenicillin, where growth remained similar to the baseline at the two lowest concentrations but decreased as the concentration increased, resulting in total growth inhibition at the highest concentration (Fig. 6). Collectively, these data characterize the growth of *P. melaninogenica* in the presence of 15 antibiotic classes, providing a comprehensive antibiotic resistance and sensitivity profile for the organism.

**Figure 6.**
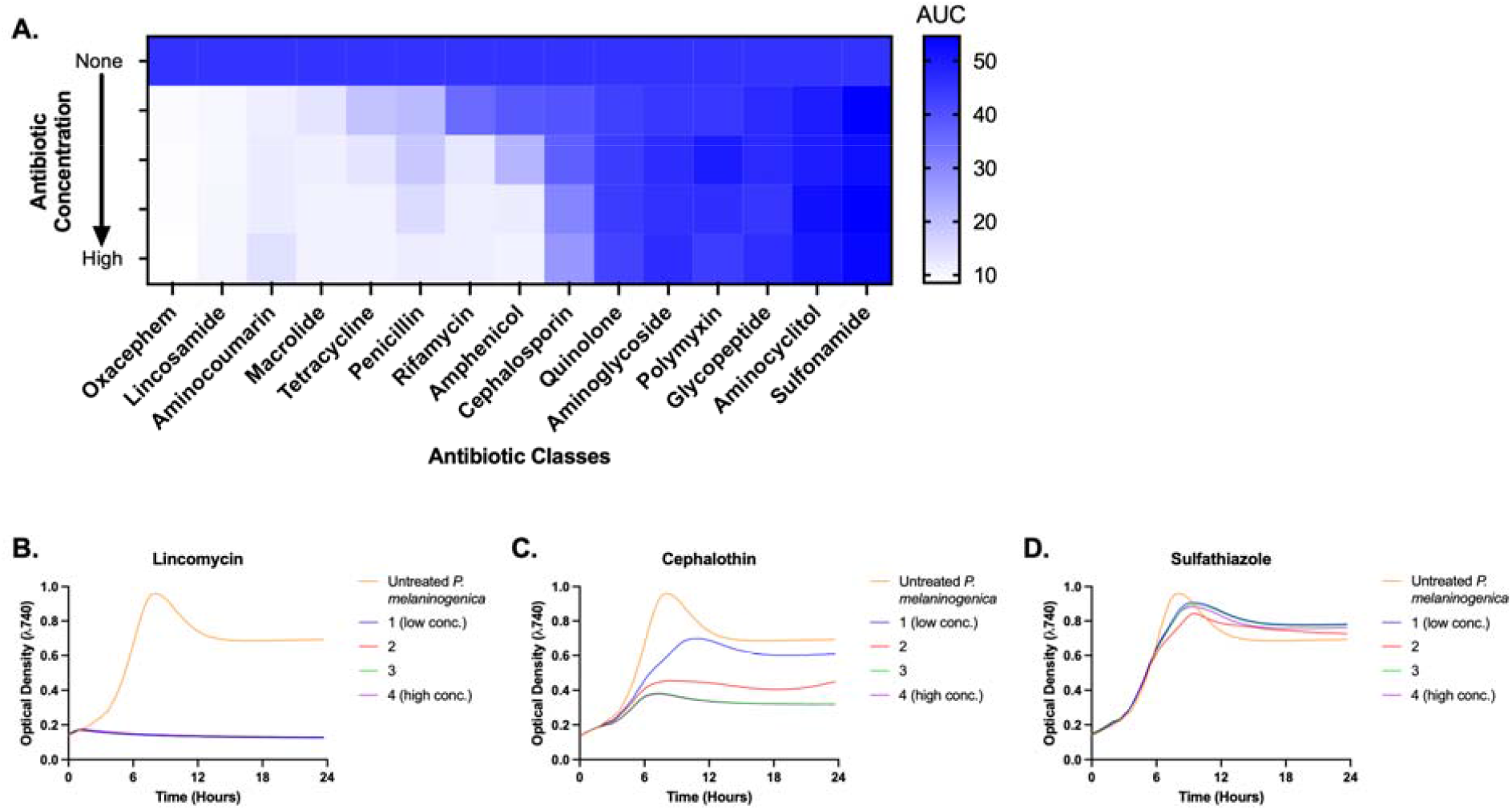
Antibiotic sensitivity profile for *P. melaninogenica* ATCC 25845. Antibiotic characterization of *P. melaninogenica* ATCC 25845. A heat map of *P. melaninogenica* responses to 15 antibiotic classes. Area under the curve (AUC) calculated from growth curves (n=3) of *P. melaninogenica* treated with 4 concentrations of each antibiotic. Representative growth curves (n=3) for sensitive (B), dose-dependent (C), and resistant (D) antibiotic groupings.

### P. melaninogenica prefers carbohydrates

To understand the metabolic needs of *P. melaninogenica*, our objective was to determine its carbon utilization profile. Because *P. melaninogenica* is present across a range of environmental niches in the respiratory tract, mapping its ability to utilize a variety of nutrients is highly relevant to understanding its colonization mechanisms and role in community dynamics. To do this, we tested *P. melaninogenica growth* on a panel of 190 carbon sources. To quantify changes in *P. melaninogenica* growth (hereby called Δ) between supplemented cultures and unsupplemented negative control cultures, we used Bacterial Basal Media (BBM). This basal media allows for detection of significant Δ when *P. melaninogenica* is supplemented. The Δ was quantified as the difference in area under the curve between the control and supplemented cultures. A Δ area under the curve of >5 was considered significant. Of the 190 carbon sources tested, 17 demonstrated a significant positive impact on *P. melaninogenica* growth (Fig. 7A). The three carbon sources with the highest +Δ were: D-maltose (+Δ= 41.70), alpha-D-glucose (+Δ= 33.37), and Lactose (+Δ= 29.09). Interestingly, all 17 of these carbon sources were carbohydrates. No carboxylic acids, amino acids, esters, alcohols, amines, fatty acids, polyols, nucleosides, peptides, or amides tested had a significant positive effect on *P. melaninogenica* growth. These data demonstrate that the carbon sources best utilized by *P. melaninogenica are* carbohydrates.

**Figure 7.**
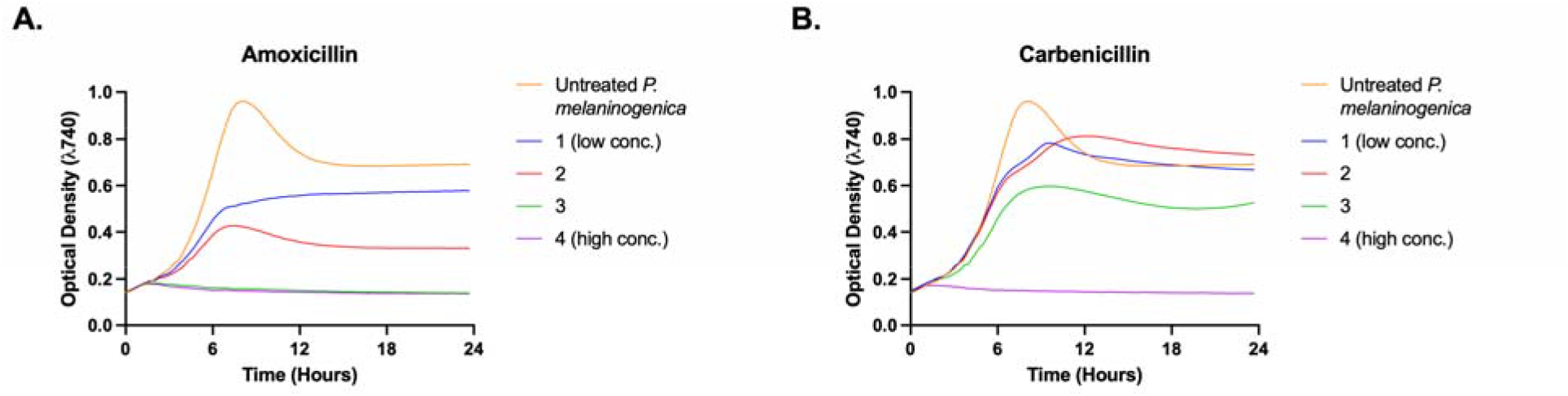
*P. melaninogenica* ATCC 25845 penicillin class outliers. *P. melaninogenica* treated with 4 concentrations of Amoxicillin **(A)** and Carbenicillin **(B)** (n=3). A dose- dependent response is seen, contrary to the complete sensitivity to the rest of the penicillin class.

**Figure 8.**
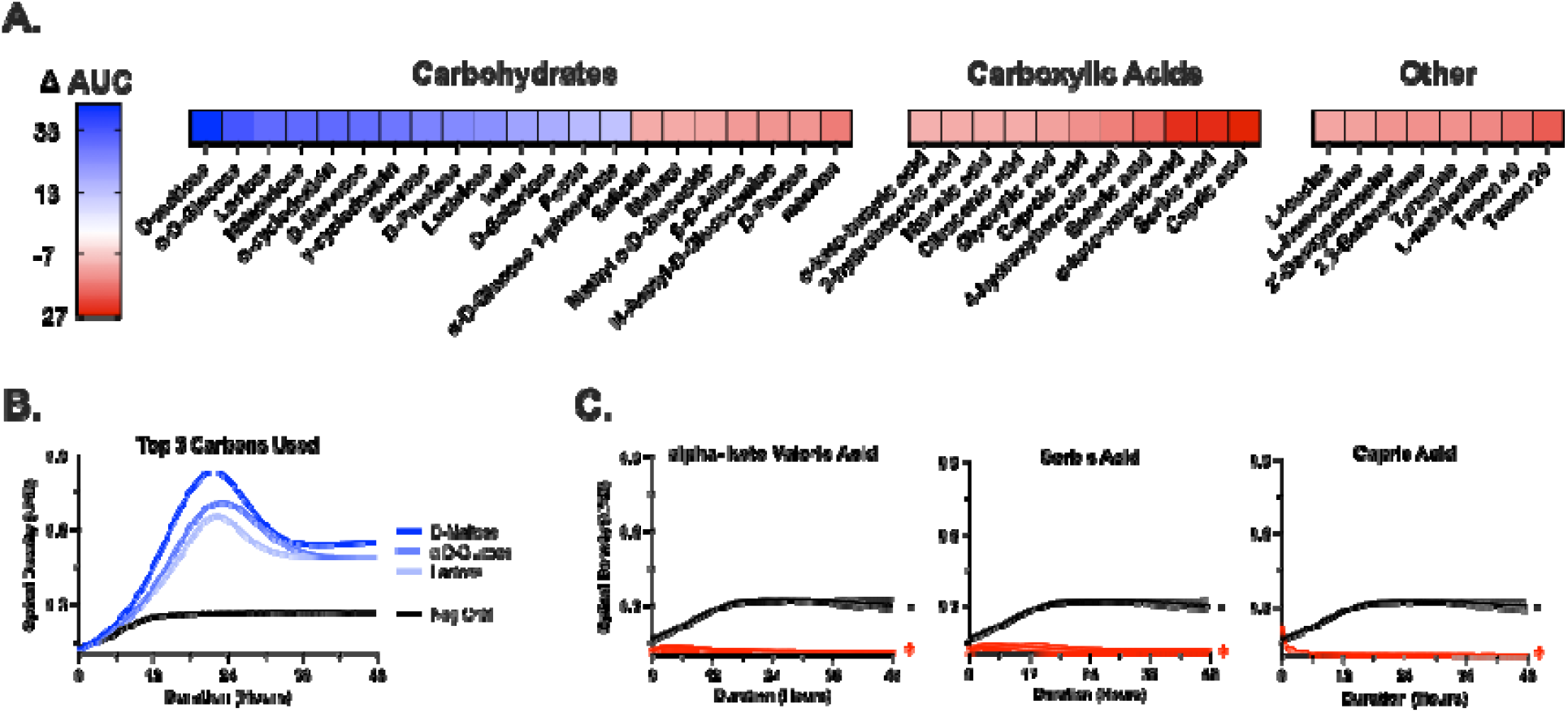
Carbon utilization profile for *P. melaninogenica* ATCC 25845. Carbon source utilization profile of *P. melaninogenica* **(A)** Screening of 190 carbon sources revealed that *P. melaninogenica* only demonstrated a significant increase in growth in the presence of carbohydrates, while some other carbons (particularly carboxylic acids), have a significant inhibitory effect on growth. **(B)** Growth curves of *P. melaninogenica* supplemented with the most utilized carbon source. **(C)** Growth curves of *P. melaninogenica* supplemented with the most growth-inhibitory carbon sources. Carbon utilization was quantified as the ΔAUC = “Change in Area Under the Curve” between the supplemented sample and non-carbon supplemented control. n = 2 biological replicates. A threshold of ΔAUC of >10 or <-10 was designated as significant.

## Methods

### P. melaninogenica reconstitution, culture conditions, and media testing

*Prevotella melaninogenica* ATCC 25845 was purchased from ATCC as a frozen lyophilized culture. Prevotella melaninogenica GAI 07411 was generously provided by collaborating personnel in Kyoto, Japan. Both bacterial strains were grown using Anaerobe Systems Brain Heart Infusion (BHI) broth (AS-872) and agar (AS-6426) in an AS-500 anaerobic chamber (0% O2, 5% H2, and 95% N2). Gram staining and Sanger sequencing confirmed the bacterium’s identity.

To measure the growth of *Prevotella melaninogenica* ATCC 25845 over time, the inoculating culture of *P. melaninogenica* was initially grown in BHI broth at 37°C under strictly anaerobic conditions (0% O2, 5% H2, and 95% N2). Inoculums (n=4) derived from this culture were cultured in BHI broth with a starting OD of 0.05 and a total volume of 100 µL. Cultures were loaded into a 96-well round-bottom plate, which was then loaded into a portable Cerillo^©^ Alto plate reader. The Cerillo^©^ measured the OD of the samples over 24 hours at 37°C under strictly anaerobic conditions.

Anaerobic CFU (colony-forming units) plating was performed on Anaerobe Systems Brain Heart Infusion (BHI) plates at hours 0, 12, and 24 of culture growth. For each culture (n=4), two technical replicates were plated. CFU/mL was plotted on a log scale.

For media testing, *P. melaninogenica* ATCC 25845 was inoculated on Anaerobe Systems BHI agar for 48 hours before plating subsequent cultures on LKV (Biolog AS-142), BRU (Biolog AS-141), PSA, BUA (Biolog 71001), TSA+, and BHI agar. For liquid media, *P. melaninogenica* ATCC 25845 was grown on TSA+ and BHI agar for 24 hours before inoculation into respective liquid media (TSB+ and BHI broth). All media testing was conducted under strictly anaerobic conditions at 37ºC prior to imaging.

### Imaging with Scanning Electron Microscopy

A 50 mL overnight culture of *P. melaninogenica* in BHI broth was grown under strictly anaerobic conditions (0% O2, 5% H2, and 95% N2) at 37°C to an OD of 2.4. From this culture, three 5 mL biological replicates were inoculated to an OD of 0.05 and grown anaerobically at 37°C. To image cells in lag, exponential, and stationary phase, the three cultures were removed and fixed at an OD of 0.24, 0.50, and 1.5, respectively. To fix samples, cultures were spun down at 3000 rpm for 5 min, resuspended in PBS, spun down at 3000 rpm for 5 min, the supernatant was decanted, and fixative was added. Fixative was >20X the pellet volume. Fixative included: 2% paraformaldehyde and 2.5% glutaraldehyde in 1 M sodium cacodylate buffer at pH 7.4. The solution was gently homogenized and incubated at room temperature for 2 hours prior to storage at 4°C. All steps were performed in a strictly anaerobic environment prior to SEM imaging. 130 cells per condition were randomly selected for quantification via ImageJ.

### Assaying Antibiotic Susceptibility

Antibiotic susceptibility assays were performed using the Biolog Phenotype MicroArray (PM) bacterial chemical sensitivity panels. The chemical sensitivity PM panels consist of two pre-configured 96-well plates (PM11C and PM12B) that contain four concentrations of each antibiotic or chemical lyophilized in individual wells, enabling simultaneous dose-response assessment of 48 antibiotics and chemicals. The untreated control condition of *P. melaninogenica* was plated in an empty Biolog 96-well plate. Several additional antibiotics - doxycycline, moxalactam, and ampicillin - and their untreated positive control condition of *P. melaninogenica* were plated in an additional 96-well plate. The concentration range for these antibiotics was selected based on commonly used laboratory concentrations and their respective minimum inhibitory concentration (MIC). *P. melaninogenica* was initially grown overnight in 5 mL of supplemented Tryptic Soy Broth (TSB+) at 37°C under strictly anaerobic conditions (0% O2, 5% H2, and 95% N2). New TSB+ cultures with a starting optical density (OD) of 0.05 were prepared from this inoculating culture and added to each plate at 150 µL per well. To maintain anaerobic integrity outside the chamber, plates were sealed with SealPlate (Excel Scientific) using firm friction to ensure an airtight barrier. Kinetic growth was then monitored in an ODIN™ plate reader, with OD measurements recorded at 740 nm every 20 minutes for 24 hours. Three replicates of each panel were run and averaged to generate growth curves and area under the curve (AUC) values for each antibiotic. To generate antibiotic sensitivity groupings, AUC values of all the antibiotics within a class were averaged for each concentration group.

### Assaying Carbon Utilization

Carbon utilization was assessed using the Biolog Phenotype MicroArray (PM) system. This platform utilizes two pre-configured 96-well plates (PM1 and PM2) containing 190 unique lyophilized carbon sources, with well A1 serving as a negative control. *P. melaninogenica* ATCC 25845 was grown overnight in Brain Heart Infusion broth, pelleted by centrifugation, and resuspended in Bacterial Basal Media (BBM). The culture was adjusted to an optical density (OD) of 3.00 to achieve a final well concentration of 1.66% and a starting OD of 0.05. For each well, 2.33 µL of culture was combined with 137.67 µL of BBM for a total volume of 140 µL. The plates were sealed to maintain anaerobic integrity and growth was monitored in an ODIN™ plate reader as described above. OD measurements were recorded every 20 minutes for 24 hours, and although readings were taken at 590 and 740 nm, data analysis was restricted to 740 nm readings to prevent interference from substrate absorbance and ensure accurate quantification of cell density. Two replicates were run, and AUC values averaged between the two.

## Discussion

This research provides a comprehensive phenotypic and growth characterization of *Prevotella melaninogenica* ATCC 25845, detailing its morphological plasticity, growth dynamics, and carbon growth preferences under strictly anaerobic conditions. Morphological analysis across various media types revealed consistent round cells with entire edges and raised elevation. Gram stains demonstrated Gram-negative staining of short and medium rod-shaped cells and occasional filamentous rods. For color, *P. melaninogenica* developed tan pigment with a cream center on agar containing sheep’s blood as a principal component. This included Brucella Blood Agar (BRU), Biolog Universal Growth Agar (BUA), Laked Brucella Blood Agar w/ Kanamycin and Vancomycin (LKV), and *Prevotella* Selective Agar (PSA). This striking color formation was not observed on agar without sheep’s blood; thereby, *P. melaninogenica* grown on Brain Heart Infusion (BHI) and supplemented Tryptic Soy Agar (TSA+) exhibited opaque, cream-colored cells. Pigment production by *Prevotella melaninogenica* on blood agar plates has been observed since its description by William and Wherry in 1921 (20,2). Subsequent morphological examinations have noted its distinct grey/tan pigment on blood agar (21), but not all *P. melaninogenica* strains produce pigment; researchers have documented cases of colorless mutants that did not pigment on blood agar or laked blood agar (22). Therefore, the presence of pigment in some strains, but not others, raises questions about its role in maintaining *P. melaninogenica* growth and its origins. Several investigations have suggested protoporphyrin, not melanin, as the causative agent of age-prone pigmentation (23), as noted by its presence in other *Prevotella* species; yet many questions remain about its true role in the growth of *P. melaninogenica*.

Growth kinetic experiments established a replication rate of 3.21 hours and identified a significant phase-dependent transition where exponential-phase rods shift to smaller coccobacilli in the stationary phase as viability decreases. High-resolution imaging further uncovered an unconventional replication strategy characterized by extreme filamentation (up to 100µm) and simultaneous multi-septum division, suggesting a departure from classical binary fission. Our data demonstrate that *P. melaninogenica* has the ability to form highly filamentous rods during the lag and exponential phases but only appears as coccobacilli within the stationary phase. In bacteria such as *Escherichia coli*, filamentous rods are known to be associated with stress such as late stationary phase and oxic conditions (24). However, our results demonstrate the opposite trend. It is tempting to postulate that this unique morphology could come as a result of our second observation-the seemingly spontaneous and simultaneous division of filamentous cells at various points at once. It is possible that when *P. melaninogenica* encounters stress in the form of limited nutrients when entering stationary phase, it divides into many coccobacilli to conserve energy. This mechanism of simultaneous multiple fission is not well characterized in the literature; however, it has been found in another oral bacterium, *Corynebacterium matruchotii* (25). *C. matruchotii* elongates via tip extension at rates more than five times faster than other closely related bacterial species. It is tempting to speculate that *P. melaninogenica* may have evolved a similar replication mechanism as an adaptation to the highly variable environment of the respiratory tract. Our observations present the possibility of a novel mechanism of growth and replication in *P. melaninogenica.*

Systematic antibiotic profiling showed that while the species is highly sensitive to macrolides and lincosamides, it remains completely resistant to six antibiotic classes, including aminoglycosides and quinolones. The antibiotic classes included in this study cover several different modes of action, some of which are shared between classes and across the resistant and sensitive groups. Several classes, such as penicillin, oxacephem, cephalosporin, and glycopeptide antibiotics, function by inhibiting cross-linking of peptidoglycan in the cell wall (26-28). Of these classes, *P. melaninogenica* exhibited sensitivity at all concentrations to both penicillin and oxacephem antibiotics, while exhibiting significant resistance to cephalosporin and glycopeptide antibiotics. The response to the glycopeptide class is expected, as these antibiotics function by binding the lipid II monomer cell wall precursors, and their activity is limited to Gram-positive bacteria. Cephalosporin antibiotics bind to and inhibit penicillin-binding proteins and typically display best coverage against Gram-positives (28). Oxacephem antibiotics are structurally similar to cephalosporin antibiotics but differ at their core, where they have an oxygen atom instead of sulfur. As a result, they have increased antibacterial activity, especially against Gram-negative bacteria. Another shared mode of action (MOA) between several classes is inhibition of protein synthesis. Among the groups that share this MOA, macrolide, tetracycline, amphenicol, and lincosamide antibiotics showed complete sensitivity from *P. melaninogenica*, while the aminoglycoside and aminocyclitol classes resulted in complete resistance in *P. melaninogenica*. Both amphenicol and lincosamide antibiotics act on the 50S ribosomal subunit, with amphenicols inhibiting attachment of tRNA to the A site on the 50S ribosome, and lincosamides inhibiting peptidyltransferase and the process of peptide chain initiation (29-30). Lincosamides have been found to have strong activity against anaerobic bacteria, which is seen here in this study. Macrolide antibiotics selectively inhibit translation of certain cellular proteins by blocking the nascent peptide exit tunnel and selectively arresting the ribosome (31). Tetracycline antibiotics function by inhibiting binding of aminoacyl-tRNA to the acceptor site on the mRNA-ribosome complex (32). The effect of aminoglycoside antibiotics is unsurprising, as this class is inactive against anaerobes due to the need for active electron transport for uptake into cells (33). Aminoglycosides, along with aminocyclitol antibiotics, both bind to the 16S rRNA of the 30S ribosome, with aminocyclitols blocking translocation (34). All of these protein synthesis inhibiting classes are typically bacteriostatic, with the potential to be bactericidal at high concentrations. An additional shared mode of action across the resistant and sensitive classes is inhibition of DNA gyrase, from both aminocoumarin and quinolone antibiotics. Aminocoumarins inhibit the ATPase activity of DNA gyrase by competing with ATP for binding to the GyrB subunit (35-36). Surprisingly, they typically have limited activity against gram-negative bacteria, whereas in this study we see complete sensitivity from *P. melaninogenica*. Quinolone antibiotics inhibit both DNA gyrase and topoisomerase IV activity, which are both type II topoisomerases, enzymes essential for bacterial viability (35). The complete resistance of *P. melaninogenica* to this class is not surprising, as early generations of quinolones have a narrow spectrum of activity and resistance is common through gene mutations of efficient efflux pumps. A somewhat surprising result was the significant sensitivity observed at most concentrations to rifamycins. These antibiotics inhibit transcription by targeting RNA polymerase and typically do not exhibit activity against gram-negative bacteria due to intrinsic efflux systems that are often present (37). Polymyxin antibiotics, which insert into cell membranes, leading to their disintegration, also had an unexpected effect in this study (38). These antibiotics typically exhibit a bactericidal effect against Gram-negative bacteria, but here we observed complete resistance in *P. melaninogenica*. When comparing antibiotic profiles between *P. melaninogenica* strains, *P. melaninogenica* ATCC 25845 and *P. melaninogenica* GAI 07411, we noted several strain-dependent responses. *P. melaninogenica* GAI 07411 notably had increased resistance to several penicillin antibiotics, demonstrating full resistance to amoxicillin and carbenicillin, differing from the dose-dependent response seen with ATCC 25845. GAI 07411 also showed a dose-dependent response to polymyxin antibiotics and vancomycin, a glycopeptide, rather than full sensitivity. While ATCC 25845 demonstrated a dose-dependent response to amphenicol antibiotics, we observed complete sensitivity in GAI 07411. The differing responses of *P. melaninogenica* ATCC 25845 to antibiotic classes with similar MOAs demonstrate the need to understand a species’ complete antibiotic resistance and susceptibility profile to determine which antibiotics will be useful in a laboratory setting and to inform clinical antibiotic prescription decisions. It is further essential to understand how strain-level differences alter the behavior of a bacterium, as we see here that between strains there can be striking differences in the response to antibiotics.

Finally, carbon utilization assays testing 190 potential substrates revealed a specialized metabolic profile for *P. melaninogenica* 25845, with significant utilization restricted exclusively to carbohydrates—most notably D-maltose and α-D-glucose. Crucially, these carbon utilization results were not identical to those of other tested strains, such as *P. melaninogenica* GAI 07411; this strain-level diversity highlights the importance of strain-level testing to fully capture the metabolic capabilities of the species. Given the known abundance of *P. melaninogenica* in the starch-rich oral microbiome, our findings offer key insights into its ecological lifestyle within this niche, where exclusive reliance on carbohydrates likely indicates a preference for specific host environments. While expanding future testing to include mucins—a predominant carbon source in the lower airway—will further elucidate its metabolic capabilities across host sites, our thorough evaluation of upper airway nutrients provides the first look at the metabolic profile supporting robust colonization by this abundant commensal. Consequently, future investigations should employ comprehensive genomic and phenotypic analyses to clarify distinctions between strains and their carbon-source preferences. Meticulous assessment of a strain’s nutrient usage will support more robust research into *P. melaninogenica* within the human microbiome, uncovering critical evidence on its ecological localization, surrounding bacterial community, and oxygen tolerance and capacity.

In conclusion, this study establishes a physiological framework for *Prevotella melaninogenica* ATCC 25845, revealing a degree of biological complexity that challenges traditional models of anaerobic bacterial growth. The observation of extreme filamentation followed by simultaneous multi-septum division suggests that *P. melaninogenica* may utilize an unconventional replication strategy—potentially an evolutionary adaptation, similar to *C. matruchotii*, for rapid niche colonization within the fluctuating environment of the respiratory tract. Furthermore, the distinct transition from filamentous rods to coccobacilli during the stationary phase suggests a possible energy-conservation mechanism in which large cells divide into smaller units to survive nutrient limitation. Our comprehensive profiling reveals a highly specialized organism that maintains a narrow, carbohydrate-centric metabolic niche while exhibiting a nuanced antibiotic profile, including unexpected resistance to polymyxins and sensitivity to aminocoumarins. These findings underscore the importance of species-specific characterization in understanding how *P. melaninogenica* functions as a robust commensal and how its unique physiological plasticity may contribute to its role in community dynamics.

## Figure Legends

**Figure 1**. Gram stain of *Prevotella melaninogenica* displaying Gram-negative coccobacillus morphology. The organism was cultured on solid brain heart infusion (BHI) media for 24 h under anaerobic conditions prior to staining and imaging. Scale bar = 10μm.

**Figure 2**. Liquid culture phenotypes of *P. melaninogenica* ATCC 25845. **(A)** Growth in liquid BHI broth shows partial homogeneity with minor sedimentation. **(B)** Growth in liquid TSA broth exhibits near-total cell sedimentation. Cultures (n= 3 biological replicates per condition) were grown for 24 hrs at 37ºC under anaerobic conditions prior to imaging. Images are representative of phenotypes observed consistently across all biological replicates and batches.

**Figure 3**. Colony morphology of *P. melaninogenica* ATCC 25845 on nutrient and selective agar media. Pigmented colony morphology was observed on LKV **(A)**, BRU **(B)**, PSA **(C)**, and BUA **(D)**.Opaque cream-colored colony morphology was observed on TSA+ **(E)** and BHI **(F)**. The observed phenotypes were reproducible across independent experimental batches.

**Figure 4**. *P. melaninogenica* ATCC 25845 exhibits a 3.21-hour replication rate (*r.r.*) and significant growth phase-dependent changes in cell morphology. **(A)** Culture optical density (OD_600_) was recorded over 24 hours (n = 5 biological replicates). BHI media + *P. mel* (Blue line); BHI media – *P. mel* (Grey line). Viable cell density (CFU/mL) was quantified at 0, 12, and 24 hrs under anaerobic conditions (n = 2 technical plating replicates per biological replicate). Statistical significance for differences in cell viability (ΔCFU) was determined using an unpaired Welch’s *t*-test with a Gaussian distribution assumption. ΔCFU was statistically significant between 0–12 h (p = 0.0008) and 0–24 h (p = 0.004). **(B)** Scanning electron microscopy (SEM) images of cells harvested during lag, exponential, and stationary growth phases (12,000× magnification; scale bar = 2 µm). **(C)** Quantification of cell length (µm) in exponential versus stationary phases (n = 130 randomly selected cells per condition, quantified via ImageJ). Statistical significance for differences in cell length (ΔCell Length) was determined using an unpaired Welch’s t-test with a Gaussian distribution assumption. ΔCell Length between exponential and stationary phases was statistically significant (p < 0.0005).

**Figure 5**. Scanning electron microscopy (SEM) images of P. melaninogenica filamentous morphology and division during exponential growth phase. **(A)** Representative SEM image showing an elongated, filamentous cell. These filaments, with a length of ≥ 100 μm, are outliers to the short rod morphology found in exponential phase. Scale bar = 10μm. **(B)** Exponential-phase filamentous cells undergoing simultaneous multiple fission. Yellow arros denote sites of septal pinching. Scale bar = 10μm.

**Figure 6**. Antibiotic characterization of *P. melaninogenica* ATCC 25845. **(A)** Heat map of *P. melaninogenica* responses to 15 antibiotic classes. Area under the curve (AUC) calculated from growth curves (n=3) of *P. melaninogenica* treated with 4 concentrations of each antibiotic. Representative growth curves (n=3) for sensitive **(B)**, dose-dependent **(C)**, and resistant **(D)** antibiotic groupings.

**Figure 7**. *P. melaninogenica* treated with 4 concentrations of Amoxicillin **(A)** and Carbenicillin **(B)** (n=3). A dose-dependent response is seen, contrary to the complete sensitivity to the rest of the penicillin class.

**Figure 8.** Carbon source utilization profile of *P. melaninogenica* **(A)** Screening of 190 carbon sources revealed that *P. melaninogenica* only demonstrated a significant increase in growth in the presence of carbohydrates, while some other carbons (particularly carboxylic acids), have a significant inhibitory effect on growth. **(B)** Growth curves of *P. melaninogenica* supplemented with the most utilized carbon source. **(C)** Growth curves of *P. melaninogenica* supplemented with the most growth-inhibitory carbon sources. Carbon utilization was quantified as the ΔAUC = “Change in Area Under the Curve” between the supplemented sample and non-carbon supplemented control. n = 2 biological replicates. A threshold of ΔAUC of >10 or <-10 was designated as significant.

**Supplemental Figure 1**: *P. melaninogenica* demonstrates complete sensitivity to the majority of penicillin antibiotics. *P. melaninogenica* cultures were treated with 4 concentrations of Cloxacillin **(A)**, Penicillin G **(B)**, Ampicillin **(C)**, Nafcillin **(D)**, and Oxacillin **(E)**.

**Supplemental Figure 2:** Screening of 190 carbon sources for impact on the growth of *P. melaninogenica*. Carbon souces are categorized by chemical class: carbohydrates, carboxylic acids, peptides, amino acids, alcohols, amides, esters, nucleosides, fatty acids, and amines. Carbon utilization was quantified as the change in Area Under the Curve (“ΔAUC”) between each supplemented culture and non-carbon supplemented control. For the purpose of this analysis, an ΔAUC of >10 or <-10 was considered significant. n = 2 biological replicates / condition. Carbons significantly enhancing growth (ΔAUC of >10) are indicated by up arrows (↑), Carbons significantly inhibiting growth (ΔAUC of < -10) are indicated by down arrows (↓), and carbons producing no significant change in growth (|ΔAUC| ≤ 10) are labeled as non-significant (“ns”).

---

**Supplementary Figure 1:**
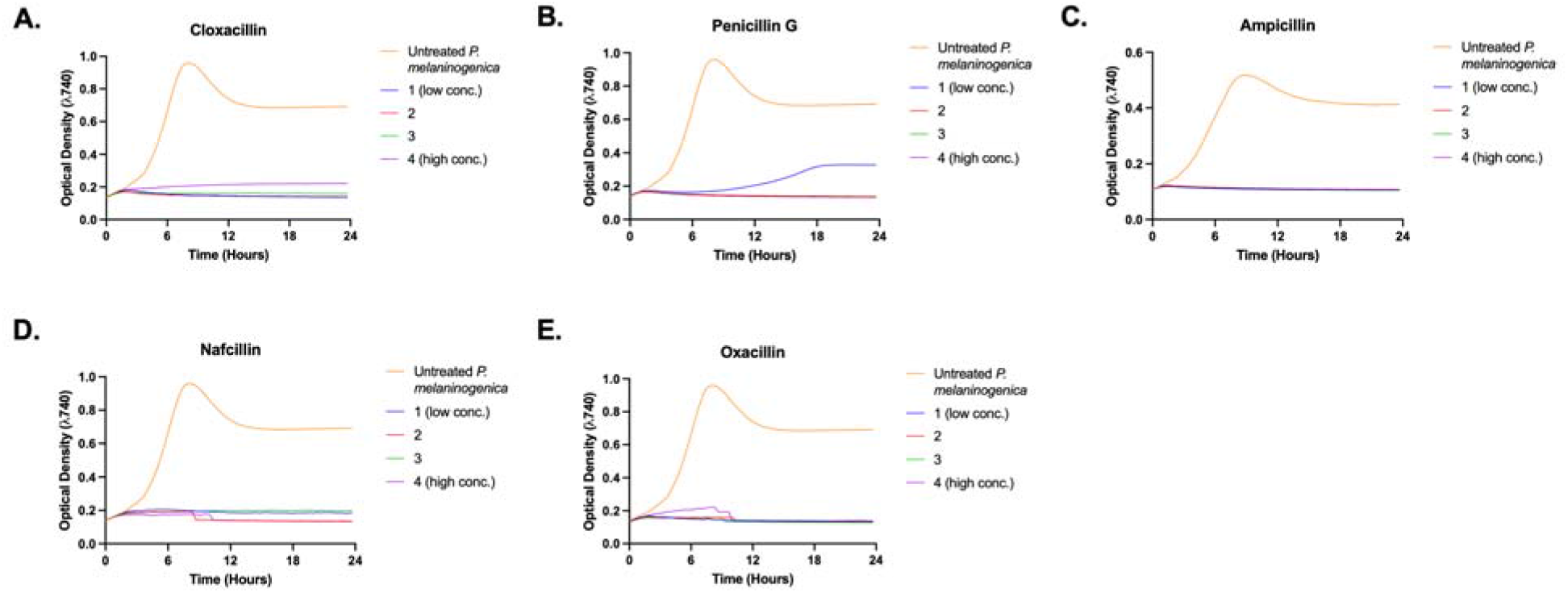
Sum of penicillin class growth curves for *P. melaninogenica* ATCC 25845. *P. melaninogenica* demonstrates complete sensitivity to the majority of penicillin antibiotics. *P. melaninogenica* cultures were treated with 4 concentrations of Cloxacillin **(A)**, Penicillin G **(B)**, Ampicillin **(C)**, Nafcillin **(D)**, and Oxacillin **(E)**.

**Supplementary Figure 2:**
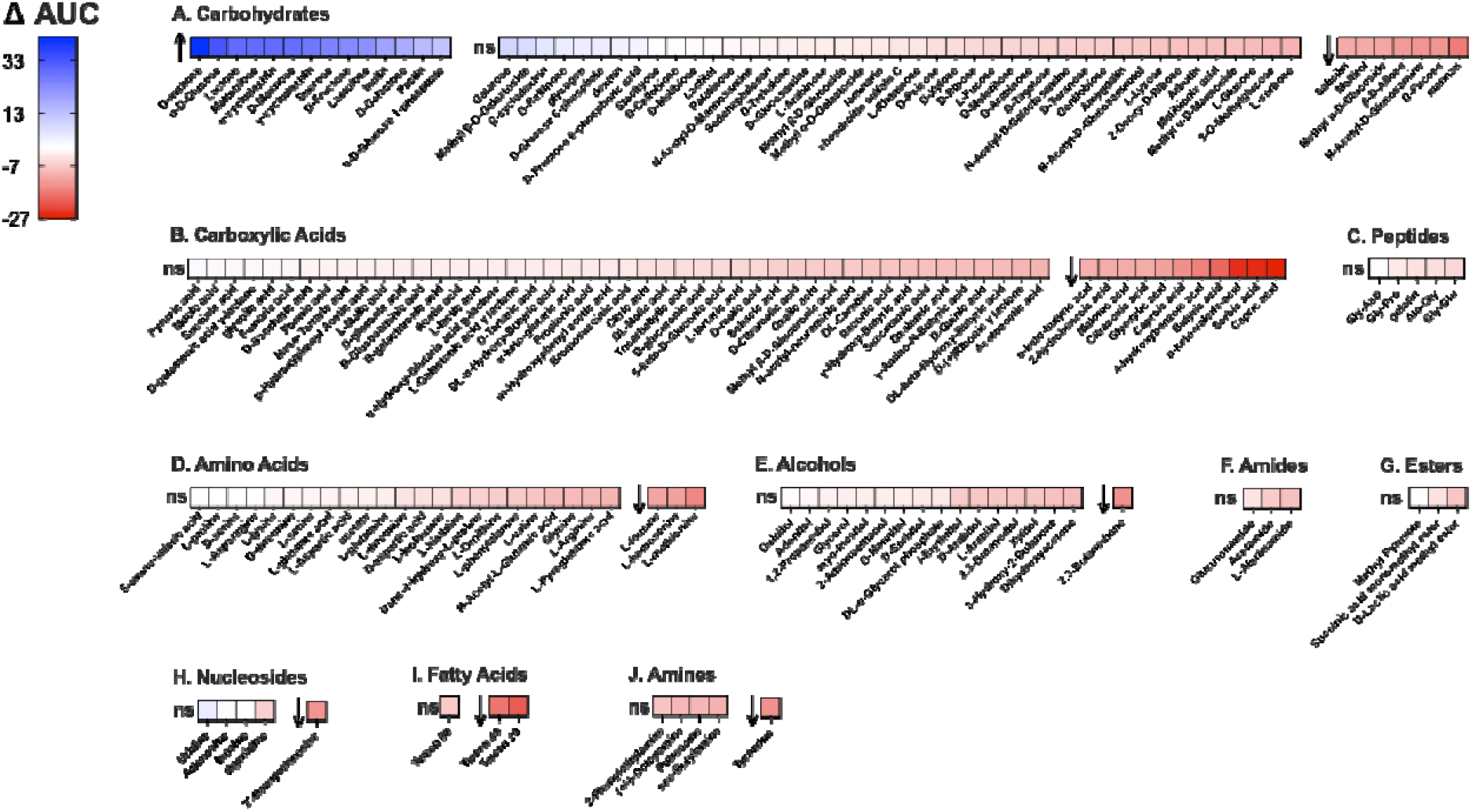
Sum of penicillin class growth curves for *P. melaninogenica* ATCC 25845. Screening of 190 carbon sources for impact on the growth of P. melaninogenica. Carbon souces are categorized by chemical class: carbohydrates, carboxylic acids, peptides, amino acids, alcohols, amides, esters, nucleosides, fatty acids, and amines. Carbon utilization was quantified as the change in Area Under the Curve (“ΔAUC”) between each supplemented culture and non-carbon supplemented control. For the purpose of this analysis, an ΔAUC of >10 or <-10 was considered significant. n = 2 biological replicates / condition. Carbons significantly enhancing growth (ΔAUC of >10) are indicated by up arrows (↑), Carbons significantly inhibiting growth (ΔAUC of < -10) are indicated by down arrows (↓), and carbons producing no significant change in growth (|ΔAUC| ≤ 10) are labeled as non-significant (“ns”).

